# DSeg: A dynamic image segmentation program to extract backbone patterns for filamentous bacteria and hyphae structures

**DOI:** 10.1101/289652

**Authors:** Hanqing Zhang, Niklas Söderholm, Linda Sandblad, Krister Wiklund, Magnus Andersson

## Abstract

**Motivation:** Quantitative image analysis of growing filamentous fungi and prokaryotes are important to detect and evaluate morphological effects of growth conditions, compounds and mutations. However, analysis of time-series image data is often limited by the ability of the algorithms to accurately segment structures that are complicated or if an organism is within a crowded population. To overcome these issues we present DSeg; an image analysis program designed to process time-series image data as well as single images to find multiple filamentous structures e.g., filamentous prokaryotes, yeasts and molds using a dynamic segmentation approach. DSeg automatically segments and analyzes objects, includes drift correction, and outputs statistical data such as persistence length, growth rate and growth direction.

**Availability and implementation:** DSeg is a free open-source program written in MATLAB. DSeg can be downloaded as a package from https://sourceforge.net/projects/dseg-software.

**Supplementary information:** Supplementary data are available at online.

## 1 Introduction and motivation

In microbiology, cell biology and biophysics, quantitative evaluation of morphological organism changes - for example, measures of persistence length, growth rate and growth direction - in time-series image data is crucial for understanding the dynamic processes of cell growth. These experiments produce large amount of data that often requires manual labor-intensive analysis. However, the development of image-processing algorithms and software can provide substantial help to users to analyze and quantify cell morphology Jones *et al.* (2008); Arganda-Carreras *et al.* (2010); Heller *et al.* (2016). Many of these analysis tools are however limited by the ability of their algorithms to extract an organisms structure from experimental images that contain noise or are of low quality. In addition, when studying growing organisms in crowded environments, organisms often grow into each other producing shape distortion and overlapping whereas long time studies often are affected by distortion, intensity variation and overlapping regions (Fig 1A). Also, regular thresholding can cause high levels of noise in the image as well as broken segments that give incorrect results (Fig 1B).

**Fig. 1.**
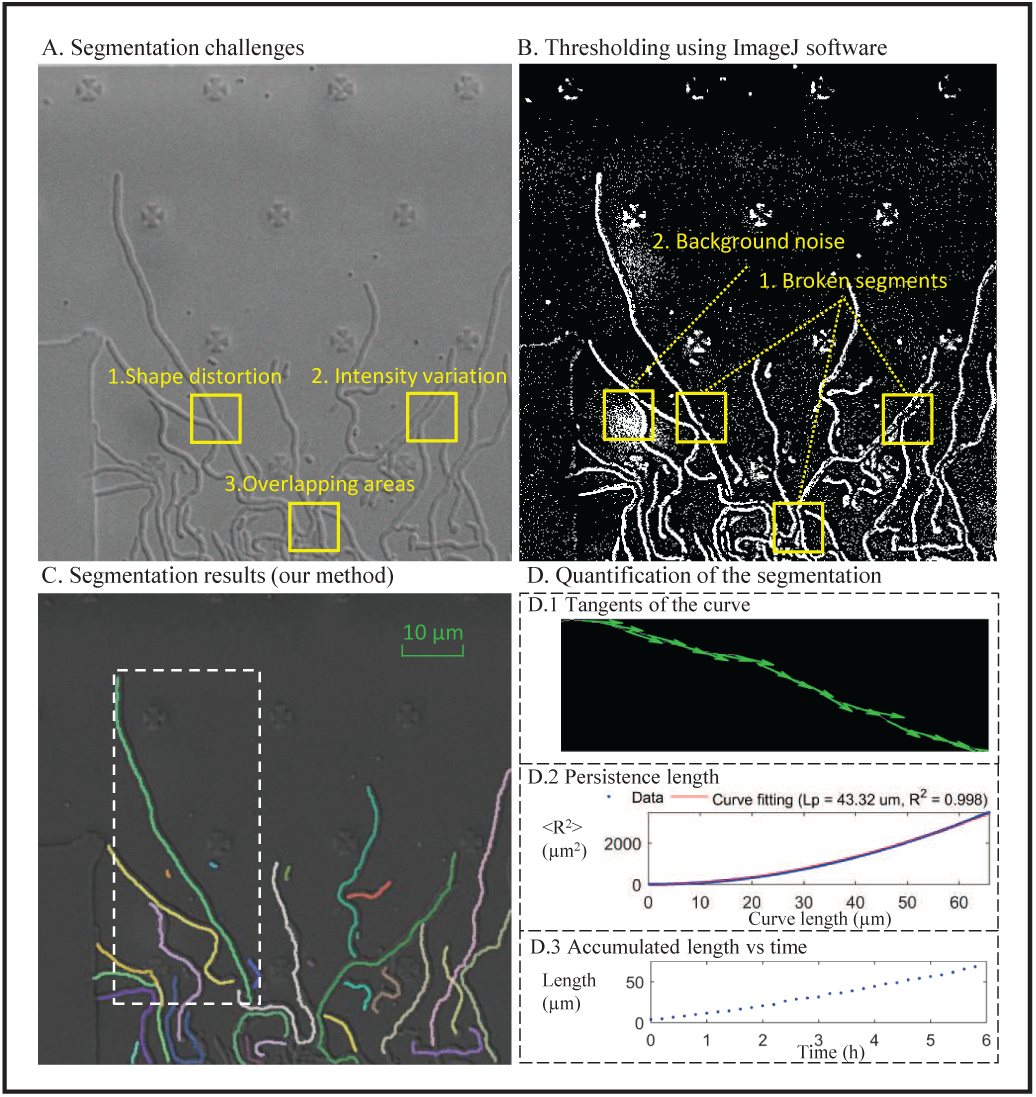
A) Three typical problematic image segmentation scenarios when extracting edges of hyphae: Shape distortion; intensity variation; overlapping areas. B) Common problems when segmenting cells using thresholding. C) Segmentation results using the dynamic segmentation algorithm in Dseg. D) Results using the segmented section in panel C (dashed box). The panels show tangents of the curve, the persistence length and the accumulated length of the bacteria under 6 h growth.

To overcome these problems and improve the accuracy of both shape analysis and quantification of structural parameters, we present an image analysis tool, DSeg, that use a specific workflow to segment static or time-lapse image data (Fig S1). We implemented a fast binary level-set based algorithm combined with constrains of size, edge intensity and growth rate for measuring organism contours and extracting patterns for analysis.

## 2 Program features and work principle

We implemented DSeg in MATLAB and provide a user-friendly interface to load, process and analyze images or time-series video data (Fig S2). We provide in the Supplementary materials an instructive user guide how to set parameter values for best performance and details about the implemented algorithms. We also provide a video that shows how to get started and how to perform an analysis.

### 2.1 Image segmentation with drift correction

After a video is loaded the user can decide whether to make a fully automatic segmentation or a semi-automatic segmentation by selecting which objects to analyze using a point-and-click approach. We recommend the latter if images are of low quality - contains pronounced image noise from intensity fluctuations or have low contrast, and if the objects under study are within a crowded environment. For the semi-automatic approach the user can set optimized parameter values for each object to segment. This is in particular useful if light conditions are fluctuating over the time scale of the experiment.

Microscope imaging experiments of growing organisms under long time scales often requires that the sample stage is moved in a matrix-like pattern to record a large field of view. Images are then captured as stacks, where each stack represent a given position of the stage for a given time. This movement, as well as the long experimental time often give rise to drifts that are clearly seen when sweeping through the stack of images. Even drifts of a few micrometer can produce significant problems if one want to automatically follow the growth pattern of an organism. To handle this, DSeg includes a drift correcting algorithm that can improve and simplify long time experiments where a large field of view is desirable.

### 2.2 Working principle of the backbone pattern extraction

To extract object contours and refining shapes using constraints of size, edge intensity and growth rate, we combined thresholding, a fast binary level-set segmentation method and a backbone structure extraction algorithm based on the framework presented in Zhang *et al.* (2017). Objects are first selected in the starting frame using the point-and-click function to find their structures from a binary image obtained using thresholding. After selecting the objects to track the automatic level-set based segmentation algorithm is initiated to track the growing pattern of the object until the end frame, see the supplementary materials file for more details.

The algorithm is robust and can handle background changes between frames as well as bacterial shape deformation. To refine the segmentation results for analysis, the user can manually segment or delete objects if the auto-detection process did not work properly. For example, these can be objects that entered from the outside of the region of interest, objects that did not grow, or objects for which the initial parameter values did not work. This process ensure correct tracking of backbone patterns with the least user intervention. When all objects have been segmented the corresponding data can be used for quantifying the objects; width, length, persistence length, growing rate, growing direction of the backbone. Since raw data is saved the user can also extract other user-specific parameters.

## 3 Demonstration and evaluation of DSeg

To verify the performance of the DSeg program we grow hyphae forming bacteria, *Streptomyces coelicolor* (*S. coelicolor*), in a fluidic chamber and imaged these using differential interference contrast and using a modified microscope in which we introduced digital holography Dahlberg *et al.* (2018); Stangner *et al.* (2017). For details about the experiment see the supplementary materials section. To record a large field of view, we programmed a motor- and piezo-stage to move in a 2×12 matrix, providing a final field-of-view of 400×1200 *µm*^2^. We sampled an image in each area every 10 minutes during a 24-hour-long growth experiment, thus generating 24 stacks of images each containing 144 images.

To dynamically segment hyphae and find specific structural parameter values, we choose the objects of interest at an early stage, that is, before the *S. coelicolor* spores started to grow. Since the bacteria grow in a highly populated area and overlapped at some regions we used the semi-automatic segmentation approach following the analysis process (Fig. 1C). We used the final segmented data to assess hyphae parameters and show in Fig. 1D examples of these, for example, tangents of the curve, persistence length and accumulated length. For representative purpose we colored each segmented hyphae differently.

We also compared the segmentation precision of DSeg with other segmentation algorithms and softwares Klein *et al.* (2012); Dimopoulos *et al.* (2014); Strisciuglio and Petkov (2017). We found that DSeq in general performed better than all other tested programs. We include a quantitative analysis of this comparison in the supplementary materials, Fig S3 and Table S1. In addition to these experiments we show as a proof of concept that DSeg can segment various images/videos. For example, we extracted structural parameters of negative stain CFA/I adhesion pili Brown *et al.* (2018), diffusion of a F-pilus Silverman and Clarke (2010), and digital in-line holographic microscopy data of *S. coelicolor* (Fig S4).

## 4 Conclusion

DSeg provides a GUI to allow for an easy and a fast evaluation procedure to assess growth related parameters of filamentous organisms such as fungi and prokaryotes etc. For example, parameter values such as persistence length, area and growth rate can be assessed directly in the program, however, since the segmented pixel values are automatically exported, other user-specific parameter values can be investigated. DSeg includes drift correction which is useful when acquiring images during long time scales.

## Supplementary materials

## 1 Requirements

We developed DSeg in MATLAB (version R2016b) using the image processing toolbox. For easy processing of video files, a graphical-user-interface (GUI) is included. DSeg is tested on Windows 7 and 10 and is made for a 64-bit operating system. We recommend a minimum of 4 GB of RAM memory and sufficient hard drive space to ensure reliable operation when running long video files. We recommend at least a 2.0+ GHz processor and to display the GUI properly the screen resolution should be 1920×1200. To run the program source code it is necessary to install MATLAB, however, to run the executable version of the program, the MATLAB Runtime R2016b can be installed without installing MATLAB. This can be downloaded at www.mathworks.com. DSeg process both images and video files and the program support a wide variety of formats, for example .png, .jpg, .tif, .avi, .mp4, .m4v, .mov.

## 2 Workflow – from experiments to analysis

Here we show the workflow of how to analyze long-term video data with DSeg using *Streptomyces coelicolor* as example (Fig S1). The procedure includes four major steps: load dataset, preprocessing data, tracking dynamic growth, and static as well as dynamic analysis of the segmented data. Note that the quality of the data is very important for reliable analysis. Thus, careful design of the experimental procedure is needed to reduce noise and optimize contrast.

**Figure S1.**
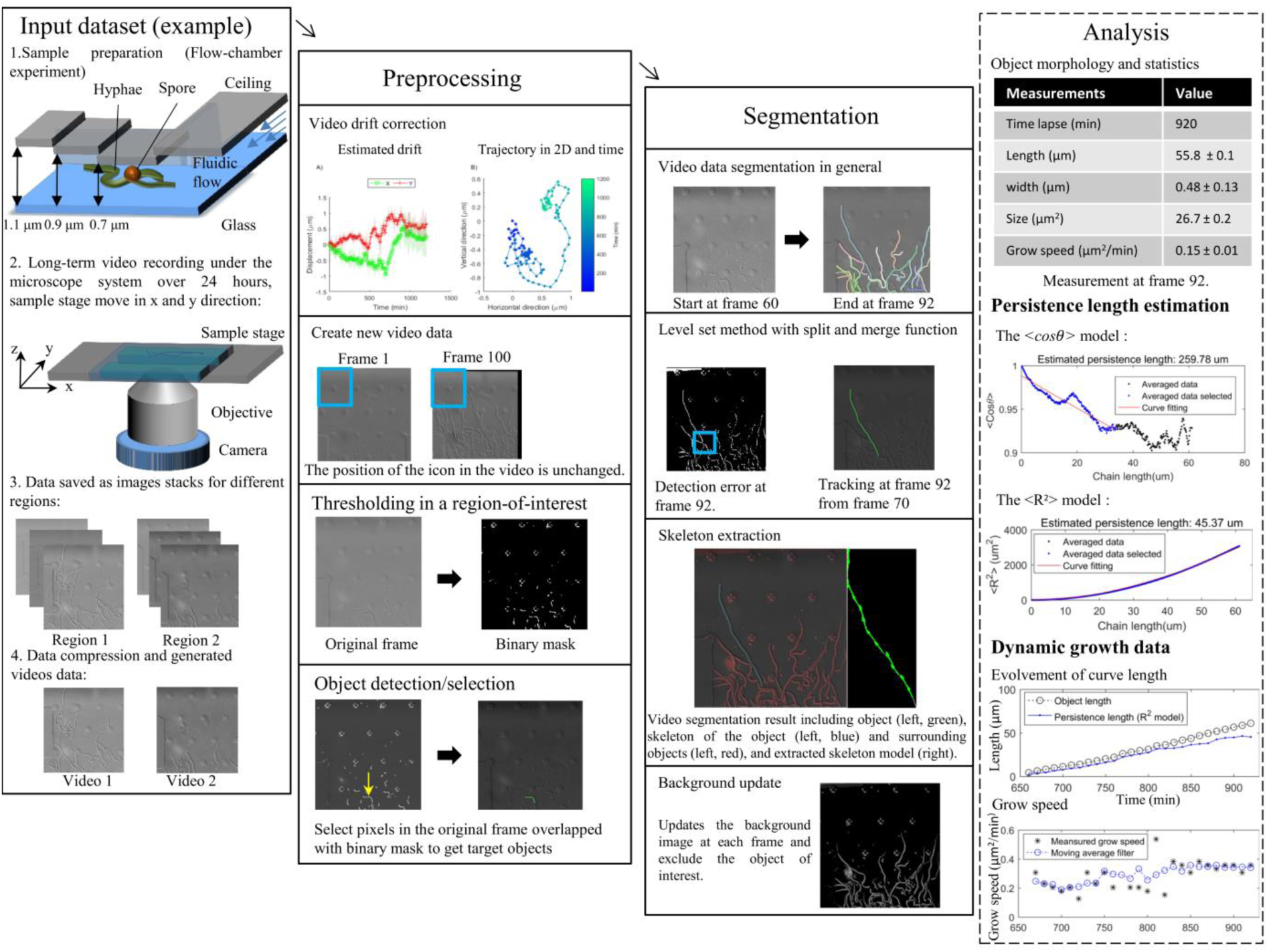
Workflow chart showing the four steps in the process from loading data to analyzing cells.

### Time lapse imaging of *S. coelicolor*

Bacteria were grown in thin microfluidic plates (CellASIC ONIX) to restrict the growth in 2D. We prepared the plates according to the manufacturer’s instructions. Spores of *S. coelicolor* M145 were introduced into the microfluidic chamber and the plate transferred to an AxioObserver.Z1 inverted microscope equipped with an incubation chamber set to 30° C. We set the flow of TSB media to 6 psi throughout the experiment. The growth was monitored using a 100× oil-immersion differential interference contrast objective with numerical aperture of 1.46. We set the exposure to 100 ms and captured images every 10 minutes for approximately 24 hours using an EMCCD camera (iXonUltra ANDOR).

To record digital holographic data of *S. coelicolor* we used a customized IX70 microscope (Olympus) modified for flow chamber and optical trapping experiments (Fällman et al. 2004; Dahlberg et al. 2018). We used a laser operating at 491 nm (Cobolt AB, Calpyso) that was run at 5 mW and to reduce parasitic reflections of the laser light we applied a rotating ground glass diffuser (Stangner et al. 2017). A custom written LabView program was used to control and translate the stage to acquire images over a large field of view.

### Video data analysis with DSeg

To analysis a video in DSeg we apply three steps. To begin with, we check for drifts in the video that can be induced by movements of the microscope when recording long video sequences, this includes for example thermal-, stage-, and objective drifts. Also, if a large field-of-view is needed the sample stage is moved in discreet steps to cover a large area. This stage motion can also induce drifts in a video since the stage is not always going back to the exact position. A drift correction functions is therefore implemented to adjust for these drifts which are detected by pixel translation in the x and y direction in images. Thus, with the drift correction function applied in the program a new video that is corrected for drifts can be made. This function also makes it possible to convert an images sequence into a video file.

To extract the objects of interest in a video file, uncorrected or corrected, a thresholding algorithm producing a binary mask is applied. This process thus detects the initial objects shape for the segmentation algorithm. Since the image quality can differ between experimental setups we offer both adaptive (Bradley and Roth 2007) and global thresholding (Otsu 1979) in which fine tuning of the detection can be applied. To further improve the selection of objects it is possible to set a region-of-interest and select object of interest manually to avoid false-positives.

After objects have been selected a dynamic segmentation process is performed, see *Segmentation* in Figure S2A. This is an automatic process and the user can change the parameters for segmentation to refine the results. Note that the program can handle broken segments, extract the skeleton shape of the object of interest and differentiate the target from surroundings objects.

To analyze the segmented data, we include tools to analyze both one image and a time-series image data file. The analysis provides basic morphological features of the segmented objects such as length, width and area size. In addition, growth velocity and direction can be extracted when analyzing time-series video data. Also, an estimation of persistence length based on the backbone pattern of the object can be measured using two approaches, one suitable for very curl objects (persistence length much shorter than the contour length) and one suitable for straight objects (persistence length similar to the contour length). All results are saved into .mat files automatically and can be export to .xlsx files for further analysis. Below we show how to perform such analysis using the program and we show the main functions to extract backbone patterns for filamentous structures.

## 3 Quick start

Screenshots of the graphical user interface (GUI) implemented in MATLAB are shown in Figure S2. In the main interface the file input-output functionalities and parameters can be set. To conduct an analysis using DSeg please follow the procedure below. We also suggest watching the short instruction video showing functionalities and how to run the program: https://youtu.be/qMbM0shkk7A

**Figure S2.**
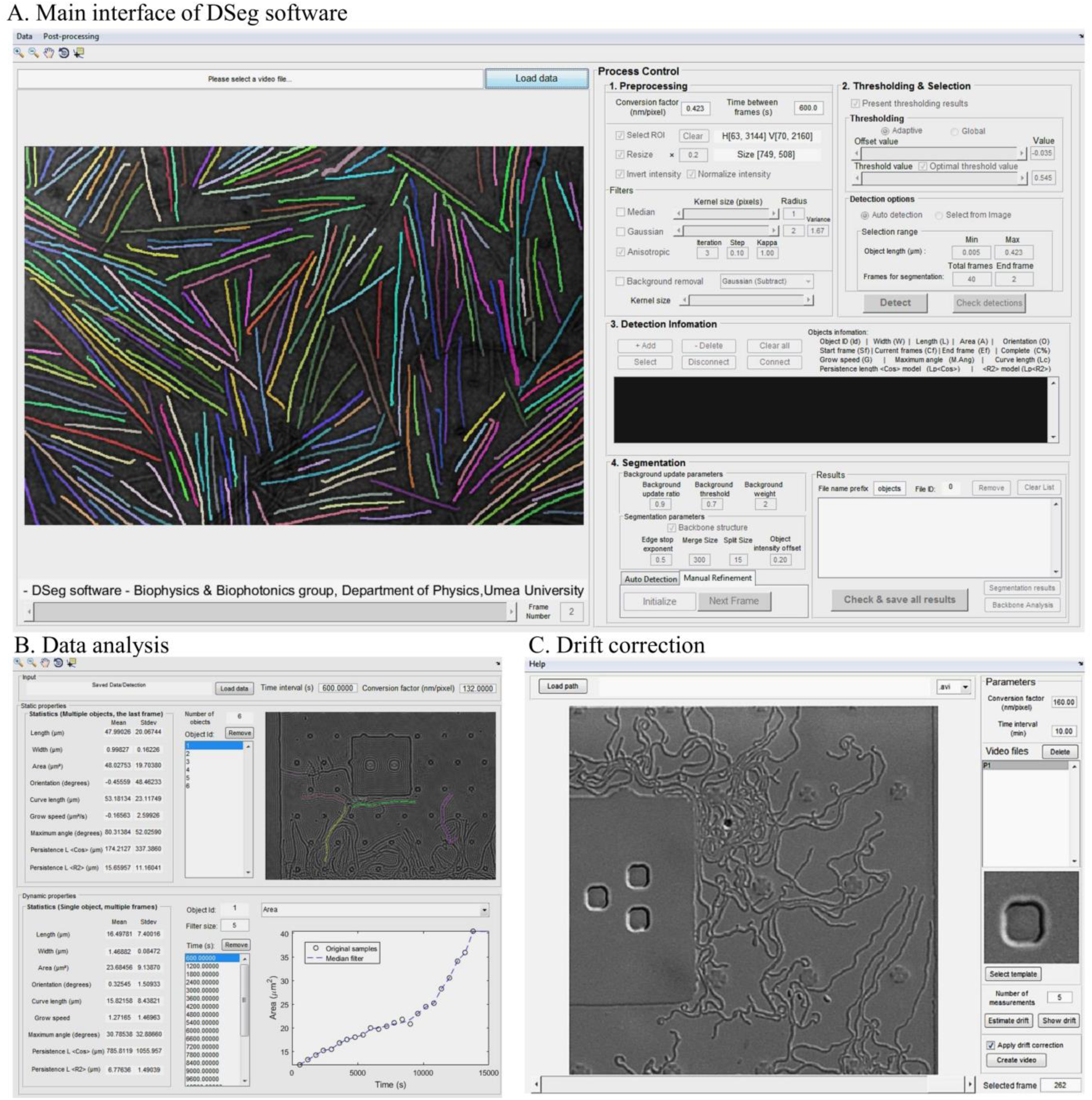
Screenshot of the main-, data analysis and drift correction interfaces. In the main interface the user controls the preprocessing filters and a function to apply thresholding to images. In addition, the detection information provides object shape properties and the status for segmentation process. The data analysis contains the statistics of both static and dynamic properties of object shape. The drift correction interface allows the user to check for drifts in the data and create a new video file.

### Basic Procedures

1. Unpack the .zip file to a folder and open MATLAB. Run DSeg.m in MATLAB to load the main interface.
2. Load a sample video or an image by pressing *‘Load data’* in the front panel, (Fig S2A) and select the video. Once the video is loaded, the file path, name and the first frame of the video will appear in the GUI.
3. In the ‘*Preprocessing*’ panel, first set the pixel-to-nm conversion factor and time between frames according to the sampling rate of the camera. In the software options: to determine a region-of-interest click ‘*Select ROI’*, to scale the video select ‘*Resize*’, to change the intensity features use ‘*Invert Intensity*’ or ‘*Normalize Intensity*’. In addition, different filters can be applied using functions in the ‘*Filters*’ panel. Furthermore, it is also possible to apply a background subtraction function.
4. To show results in the display from the thresholding function check the ‘*Present thresholding results*’. The original image will be covered by a binary mask as a result of the thresholding. ‘*Adaptive thresholding*’ is selected by default. It is possible to change the ‘*offset value*’ slider in the ‘*Thresholding method panel*’ to optimize the thresholding so object shapes are as clear as possible.
5. To select a starting frame drag the slider located under the image display or input a number to the ‘*Frame number*’ directly. The frames for segmentation can be set in ‘*Num of frames*’, or simply input the ‘*End of Frame*’ so that the software will calculate the number of frames. The ‘*Select from Image*’ option is chosen by default and the user can press ‘*Detect*’ to find all objects using a point-and-click approach. The point-and-click function is activated in the display of the GUI after the ‘*Detect*’ button is pressed. To select objects of interest, single click on an object overlapped with thresholding mask in the display, and double click on the object if it is the last object to be analyzed. If the *‘Auto detection*’ option is chosen for ‘*Detect*’ button then all objects within the range of object length are selected.
6. All selected objects will appear in the list under the ‘*Detection Information*’ panel and from here, the user can check if all the object of interest is selected correctly. If you click on a line in the list the segmentation results of the selected object will appear in the image display as a colored mask. Press ‘*Add*’ to add new object using the point-and-click function or press ‘*Delete*’ to remove one of the selected object from the list. The user can also fine tune the initial object shape using ‘*Disconnect*’ and ‘*Connect*’ button
7. To automatically segment and analyze all objects for the selected frames in a video, click ‘*Batch processing*’ in the ‘*Auto detection*’ tab. The user can also segment each object step-by-step by pressing ‘*Next frame*’ in the ‘*Manual refinement*’ tab, or reset the segmentation result of the object using the ‘*Initialize*’ button. Segmentation parameters in the ‘*Parameters for segmentation*’ panel can be changed to get optimal results. We explain in detail in the Algorithm section how each parameter affects the segmentation results.
8. All results are saved as .mat files in a path ‘*/Saved Data/Detection/*’ under the working directory by default with file names given by the texts in ‘*File name prefix*’ and the id number assigned to each object. Press ‘Check & save all results’ to load all results from files, update the latest results and save them to the files. The segmentation result of a selected object will present in the display by pressing ‘Segmentation results’, and detailed backbone structure analysis of this object can be generated by pressing ‘Backbone Analysis’.

### Software parameter settings

Here we explain the functionalities and the range of values for functions used in the DSeg.

#### Invert intensity

DSeg find objects based on intensity. We define objects as the pixels with the highest intensity values compared to a background. The *‘Invert intensity*’ function can be applied to the images with dark objects in a bright background.

#### Resize

To reduce the execution time of the segmentation process the image can be resized. The program will by default set the image size to a maximum of 800 pixels in length or width when a video or image is loaded. The user can change this value manually.

#### Filters

We include three types of noise reduction filters to improve the segmentation process. To remove sharp speckle noises use the ‘*Median*’ filter for which the kernel radius value can be set from 0 to 10 pixels. The ‘*Gaussian*’ filter is a low pass filter that can smooth features in the image with its kernel radius set from 0 to 10 pixels. The variance of the Gaussian filter can be set manually where higher variance produces a smoother image. We suggest that the value of the variance is one third of the kernel diameter. The third option is anisotropic filter (Perona and Malik 1990; Gerig et al. 1992) which can smooth the image intensities differently depending on the gradient value for each pixel. The iterations for anisotropic filter are restricted from 1 to 10 where a larger value generates a smoother image. The ‘*Step*’ and ‘*Kappa*’ values range from 0 to 1 control the smoothing effect for anisotropic filter. The step value should be set low to avoid intensity artifacts in the image.

#### Background removal

To create an image with high contrast and to remove background with unevenly distributed light intensities, apply the background removal function using a background image generated by a Gaussian low pass filter. The user can either subtract or divide each frame with the background image and the Gaussian kernel radius can be set from 1 to 30 pixels.

#### Thresholding

The adaptive thresholding function in DSeg can handle unevenly distributed background intensities, while the global thresholding method is more computationally efficient in finding object shape in a region-of-interest. The ‘*Offset value*’ is added to the thresholding values from either the adaptive function or the global method with thresholding value calculate by (Otsu 1979), in order for the results to be correct. The ‘*Threshold value*’ for global thresholding can be set manually ranging from 0 to 1.

#### Segmentation parameters

In order to find the contour of each object, a dynamic segmentation process based on the level-set function is implemented in the DSeg with four parameters to refine the results. The ‘*Edge stop exponent*’ can be set from 0 to 10. This parameter restricts the segmentation area of object when it has a value close to zero. The ‘*Merge size*’ and ‘*Split size*’ must be a non-negative integer representing the pixel number. To detect a fast growing object the ‘*Merge size*’ should be a large value. To keep the structure of a static object in a long-term recording a large value for ‘*Split size*’ is needed. The ‘*Object intensity offset*’ is set from −1 to 1. A positive offset value will restrict the contour shape to pixels with strong intensities while a negative offset value include more low intensity pixels as object. Details of the parameter are explained in the Algorithm section.

#### Background objects

To reduce segmentation errors, the segmentation process not only detects the object of interest, but also finds background objects in each frame. The ‘*Background update radio*’ set the weight for background images detected in the current frame and the ones before. For example, a ratio of 0.9 indicates a previous background is weighted 0.9 and a current background image is weighted 0.1 to form the new background image. The ‘*Background threshold value*’ determines if the background signal is strong enough to change the segmentation result. This threshold is set from 0 to 1. The ‘*Background weight*’ is used to determine the backbone structure from the segmentation result. A large weight value up to 100 can be set when the backbone structure is static in the image. On the other hand, if the object moves abruptly, this value can be set to zero.

### Data analysis

After all results are saved, the user can access these files using MATLAB or using the DSeg software by opening the *‘Post-processing*’ menu and click ‘*Objects properties*’. Then an interface will appear as shown in Figure S2B presenting both the statistics of multiple objects and the dynamic properties of each object. The user can select an object of interest from the ‘*Object list*’ and check how different morphological properties of this object change with time.

### Lateral drift correction

To ensure static a background, we have implemented a drift correction algorithm in MATLAB based on a template-matching algorithm using multi-scale image pyramid generated by Gaussian filters. This function can be found in the ‘*Data*’ menu of the program. By following the procedure of drift correction a video free from lateral drift can be created. With this process, the segmentation results can be improved significantly. Instructions of the details procedure for drift correction can be found by clicking ‘*Help*’ menu in the interface as shown in Figure S2C.

## 4 Segmentation method

### Intensity thresholding and object selection

Images acquired in microscopy experiments often differ in quality due to intensity fluctuations and noise, and the objects to track are often dissimilar. To design software that can handle data from different imaging systems and objects, a flexible thresholding value is required to get a binary mask that can represent the shape of the target object. This thresholding value can automatically be determined by a global image thresholding approach, for example by using the Otsu’s method. However, results from automatic thresholding can be inaccurate due to many factors, such as uneven illumination, intensity variation of the objects, artifacts from the sample device etc. Therefore, we implemented both adaptive thresholding and global thresholding based on the Otsu’s method with a possibility for the user to set an offset value to optimize the binary mask representing the target object. Thus the user can see if the shape of the object of interested is matched by the binary mask.

This binary mask works only as pre-step for the user to select which object to be analyzed. After finding the proper thresholding value, the user selects seed points, which are the objects to segment dynamically. For this step the binary mask evolves only for the selected object so that the algorithm can find the growing-pattern locally with minimum risk of getting false-positive detections from the neighboring objects.

### Segmentation algorithm

Based on the initial binary mask of the selected object, the algorithm will then evolve the contour of the object to find its edges and thereby track the changes of object shape in each frame. This procedure is automatic and the workflow for this part of the algorithm is described in detail below.

#### Edge stop and level-set functions

To find the shape of a selected object we use the level-set method where the object’s estimated contour is optimized using image gradient information. We define *I*(*x,y*) to represent the intensity of an image where *x* and *y* are coordinates along the horizontal and vertical axis, respectively. We first calculate the gradients *g*_*x*_(*x,y*) and *g*_*y*_(*x,y*) of *I*(*x,y*) in *x* and *y* directions using the Sobel gradient operator (Sobel and Feldman 1968). To smooth edges we apply a mean filter to calculate edge responses *G*_*x*_(*x,y*) and *G*_*x*_(*x,y*) and the corresponding magnitude,

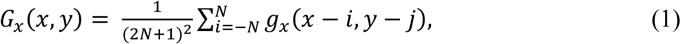

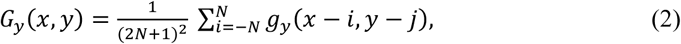

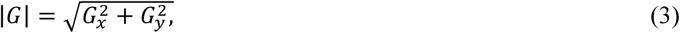

where *N* represents the radius of the mean filter kernel.

To use the gradient magnitude information in a level-set approach we define the normalized gradient, *G*_*N*_, to be the edge response magnitude, *|G|*, divided by its maximum value and we define an edge stop function by,

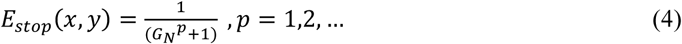

This edge stop function is used within the level-set method to restrict the evolvement of the level-set function and thereby optimize the shape estimate. Furthermore, we define the level-set function values larger than 0 to represent the foreground and values smaller than 0 to be the background so that an object can be represented as foreground surrounded by the background. When evolving the level-set function *ϕ*_*i*_(*x,y*), an implicit representation of a planar closed curve is created at zero level, the curve corresponds to *ϕ*_*i*_(*x,y*) = 0, and the changes of the curve shape is described by,

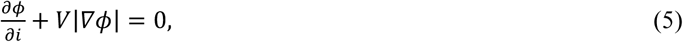

where *V* controls the evolving speed of the curve. For a binary implementation of the level-set method *V* can be represented by a sign function *F*_*s*_(*x,y*) multiplied by the edge stop function, equation 4, as (Zhang et al. 2010),

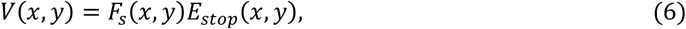

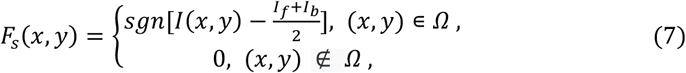

where *sgn(a)* calculates the sign of the value *a* and produce values of −1 or 1 accordingly, *Ω* represents a two dimensional region of both foreground *Ω*_*f*_ and local background around the object *Ω*_*b*_ with their averaged intensity value defined by *I*_*f*_ and *I*_*b*_, respectively. To find a complete shape of the object a good estimate of the region *Ω* is crucial. The initial estimate of *Ω* is found by using a binary image *B*(*x,y*) which consists of two parts, the foreground image *B*_*f*_ (*x,y*) and the background image *B*_*b*_(*x,y*). The foreground image *B*_*f*_ (*x,y*) is generated from the above mentioned thresholding and object selection approach. The foreground region *Ω*_*f*_ is then created by using pixels in *B*_*f*_ (*x,y*) with values equal to one. In this image we cluster connected components of 1:s using a connectivity of 8 and use the seed point to exclude components that are not in connection with the object of interest. The image *B*_*f*_ (*x,y*) contains after this process only the object of interest. The intensity *I*_*f*_ is determined by fitting a normal distribution model to the histogram of *I*(*x,y*) in *Ω*_*f*_ and determining its expectation value.

We generate the local background of the object of interest *B*_*b*_(*x,y*) by applying morphological operators, dilation and erosion, to *B*_*f*_(*x,y*) using a binary mask *K* of a typical size 5×5. The dilation and erosion process can be described in a formal way using the notation,

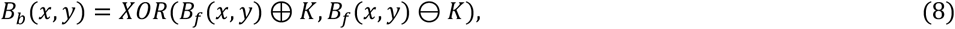

where the operator *XOR(A,B)* calculates the logic exclusive OR operation on the matrix *A* and *B* pixel by pixel, and ⊕ and erosion ⊖ are the dilation and erosion operators, respectively. The pixels with value 1 in *B*_*b*_(*x,y*) represents the local background region *Ω*_*b*_ of the object and the averaged intensity within this region from *I*(*x,y*) is *I*_*b*_. Finally, the value of *B*(*x,y*) is 1 if either *B*_*f*_ (*x,y*) or *B*_*b*_(*x,y*) is 1, so that the region *Ω* covers both *Ω*_*f*_ and *Ω*_*b*_. The binary images for foreground and background are evaluated every time when a change in the level-set function occurs. Such change corresponds to switching pixels in the contour of the object from foreground to background, or vice versa. The update of the level-set function is described by the following discrete form of equation 5,

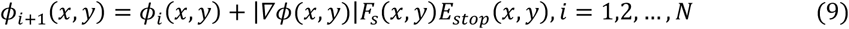

where *i* represents the level-set function iteration number and *N* defines the maximum iteration number. The initial level-set state is obtained from the *B*_*f*_ (*x,y*). In equation 9, the change of the level-set function during one iteration is applied to the contour of the object of interest according to *∇ϕ(x,y)*, controlled by the sign function *F*_*s*_ and the edge stop function *E*_*stop*_. The former one switches pixels from foreground to the local background or vice versa, and the latter one restricts the evolvement of the object contour. According to the geodesic active contour approach (Caselles, Vicent and Kimmel, Ron and Sapiro 1997) irregularities must be controlled during the evolution process. For the binary level-set method, this process is performed by using a Gaussian filter applied to the function *ϕ*_*i+1*_.

#### Split and merge operations on objects

For better accuracy in the segmentation process it is important to design an algorithm that can adapt to the features of an object and restrict the evolvement of the level-set. By visual inspection of several experimental data sets we made the following two simplifying assumptions: 1. Objects are relatively static and 2. The growing area found from two consecutive frames is small compared to the total object size. We thus only track objects that move slowly, but we allow large shape deformation. Therefore, when updating the level-set function from *ϕ*_*i*_ to *ϕ*_*i+1*_, we analyze the object region using both *ϕ*_*i*_ and *ϕ*_*i+1*_ to stop splitting one large object into small segments, and determining if newly detected objects should split or merge. We explain the details of these splitting and merging conditions below.

The condition for splitting objects is determined by comparing the foreground binary image *B*_*f*_ (*x,y*) before the level-set function evolvement with the image after one iteration 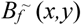. We calculate the size of lost area represented by subtracting *B*_*f*_ (*x,y*) with 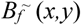 and compare the results with a threshold value using,

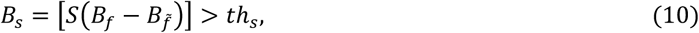

where *S(M)* calculates the area size of connected components, 1:s, in a binary matrix *M* and use this size to label each pixel in *M* accordingly. The binary image *B*_*s*_(*x,y*) represents objects which can be separated according to a threshold value *th*_*s*_. This condition stops evolvement in the level-set to avoid breaking long objects into two or more parts. In practice, we first check all pixels with the condition 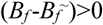 to determine if the object is shrinking. If it is true for some pixels, then the corresponding pixels in *B*_*s*_(*x,y*) are removed. It is worth noting that this restriction only applies to pixels with their area size larger than *th*_*s*_, while small objects can still be kept during the level-set evolvement.

A similar approach is used to merge two separate objects. In this case, we determine if the growing area is smaller than a threshold value *th*_*m*_ as the maximum growing rate with the condition,

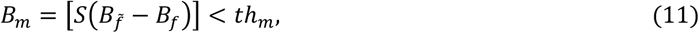

In the end, the binary image *B*_*s*_(*x,y*) and *B*_*m*_(*x,y*) from the split and merge conditions, respectively, are added to 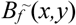. Then, as a pre-step before next level-set iteration using equation 9, we update the *ϕ* values to 1 for all 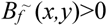 and to −1 for 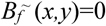.

The evolvement of the level-set function with split and merge conditions will continue until the iteration *i* reach *N* or a stop condition is fulfilled. It is important to set the stop condition objectively to avoid bias in the shape quantification process. In practice the stop condition is controlled using two criteria. The first criterion contains two conditions; the first compares pixel-by-pixel difference between *B*_*f*_ (*x,y*) and 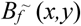 to see if the number of different pixels is smaller than a threshold number, indicating the level-set evolvement has reached a stable state. The second condition is to check if the absolute pixel difference in two consecutive iterations has the same value indicating that the error cannot be reduced further. When these two conditions are fulfilled, a second criterion is applied. This criterion update the level-set function using equation 9 with a slightly different sign function comparing to equation 7,

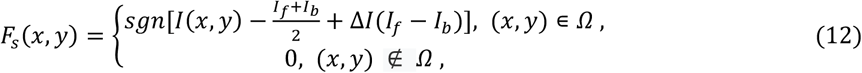

where Δ*I* is a small value in the range −1 to 1. Equation 9 with a positive Δ*I* give us the level-set function *ϕ*_Δ*I*_, and a negative Δ*I* gives *ϕ*_*-*Δ*I*_. From these level-set functions we can get foreground and background images as in the abovementioned sections. In addition, we can evaluate the intensity variance of foreground and background objects using,

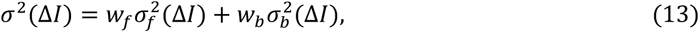

where *w*_*f*_ and *w*_*b*_ are weights calculated using the number of pixel 1:s in foreground image *B*_*f*_ (*x,y*) and local background image *B*_*b*_ (*x,y*), respectively; the 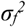 and 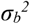 are calculated using non-zero values in *I*(*x,y*)*B*_*f*_ (*x,y*) and *I*(*x,y*)*B*_*b*_ (*x,y*), respectively. We compare the variance values to determine if both *σ*^*2*^(Δ*I*) and *σ*^*2*^(-Δ*I*) are larger than *σ*^*2*^(*0*). If this condition is fulfilled, the segmentation reaches a local optimal condition and therefore stops the level-set evolvement. Otherwise the level-set function *ϕ*_*i+1*_(*x,y*) is updated with either *ϕ*_Δ*I*_ or *ϕ*_*-*Δ*I*_ depending on which one has the smallest variance value.

#### Combining level-set with thresholding

To make a robust and accurate segmentation algorithm we implement the binary level-set method in combination with thresholding. The level-set method is a robust approach when segmenting changes of object shape over time in the presence of drifts in images. The level-set method is however, a computational expensive process and therefore needs to be optimized to be efficient. To design an algorithm efficient in finding filamentous shape, we assign the last level-set result *ϕ*_*i*_(*x,y,t*) at frame *t* directly to the next frame as the initial level-set function *ϕ*_*0*_(*x,y,t*+1). In addition, since the growing area is mainly close to the end of a filamentous structure the level-set method requires a large number of iterations to find the object shape. To reduce the number of iterations we apply a coarse estimation of the object shape using a simple global thresholding applied before the level-set evolvement.

To combine thresholding with the level-set method we define the coarse foreground of the image *B*_*cf*_ by,

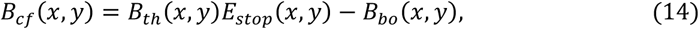

where *B*_*th*_ is a binary image from any thresholding method, *B*_*bo*_ is a binary image of background objects excluding the target object. Details of the *B*_*bo*_ are explained in the following section. The *B*_*cf*_ is compared to the foreground image *B*_*f*_ generated from *ϕ*_*0*_(*x,y,t*+1). We exclude objects in *B*_*cf*_ which are not spatially connected to the objects in *B*_*f*_ based on the rules for connectivity of 8 neighboring pixels. We set pixels values to zero for all the excluded objects and the remaining pixel values in *B*_*cf*_ are 1s. Eventually the update of the level-set function *ϕ*_*0*_(*x,y,t*+1) is completed by adding *B*_*f*_ with *B*_*cf*_ and get the objects shape under the condition *B*_*f*_ (*x,y*)>0. The growing areas are region where (*B*_*cf*_ -*B*_*f*_)>0. These growing areas are checked with the abovementioned merge conditions.

#### Background objects update

To distinguish objects of interest from other objects and the background, we use the information of background objects to avoid the level-set method to evolve an error region and apply an adaptive background update method based on the assumption that background pixels do not change abruptly in video, only small amount of movement in foreground pixels is expected. Most of the experiments related to objects growth are conducted in static background and the objects’ motions are relatively static. We get the initial *B*_*bo*_(*x,y*,0) as background objects by applying thresholding to get *B*_*th*_ and subtract *B*_*th*_ *with* the *B*_*f*_ (*x, y*). The update of background object mask *B*_*bo*_ is conducted as the following,

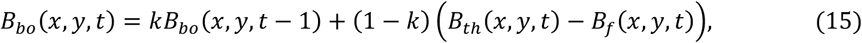

where *k* is the threshold value given as the amount of weight in the previous detection. In our case, a pixel that belongs to the background object can have a high *k* value so that the background can remain as background. In the region of target object, the *k* value also set high to keep the detection free from noises in the background. On the other hand, if an overlap is detected between the previous background object region and current object region of interest, a lower *k* value can be applied. This is due to the fact that some newly developed areas of the object can have variations in intensities which lead to the segmentation results with broken parts in the object of interest. These small parts can easily be detected as false-positive background object and get excluded from the object. Therefore the value *k* is adaptively updated with,

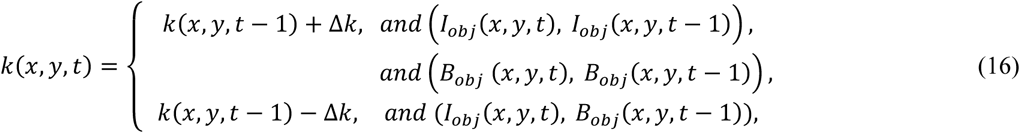

where Δ*k* is the step size, the conditions for updating *k* is implemented using logic and operator to check each pixels. In practice *k(x,y)* is assigned with initial value and the range of value is limited from 0.5 to 0.99, so that the background mask *B*_*obj*_ is dependent on at least two frames up to the 100 last frames.

## 5 Data analysis

DSeg provides shape information of objects, such as persistence length, object length, etc. This section shows how these parameter values are obtained.

### Skeletonized objects

DSeg derive the skeletonized object from the binary mask as a direct result from the segmentation process. For this we use the morphological thinning functions in MATLAB (Lam et al. 1992). This function produces a binary image that contains connected lines, in which each line is represented as one-pixel thick lines. This function allows for branching analysis as well as close loops. It is also possible to use the morphological skeleton function in MATLAB, however, the thinning function produce less branches along the contour of a segmented object making the results more accurate and easier for the analysis.

### Extracting backbone patterns

We define the longest path in the topological structure of the skeletonized object as the backbone pattern. To find the longest path, we first consider each end of the skeletonized object as a starting point, and find all the joints from the skeletonized mask. Between two joints, two starting points, or one joint and one starting point there can exist a line defined as path segment for which the algorithm can find a combination of paths that contributes to the longest distance. For each of the path segment, the distance value is the length in pixels multiplied by a positive weighted mask from the history of detected paths. We assume a simple path problem where no path can be repeated, and the solution to this problem is implemented using the extract algorithm, where all possible permutations are searched to find the solution. Since the start and end of the path must be one of the skeleton ends, and the permutations can only be selected from paths, there exist a limited number of pathways, that is significantly less than in a longest path problem. The Pseudocode of our path detection algorithm is therefore given as;

Define a node as each path segment. Find all nodes in order to a group ***N*** and label each ***n*** with its order number to

***n.id*** and calculate the properties of length ***n.len*** and neighboring nodes list ***n.list***.

~~~
**For** each staring point relate to a path segment ***n*’** in ***N***,
   Create a copy of ***N*** as ***N*’**,
   Set a check list ***L*** initialized with the label of ***n*’**,
  **While** (***L*** is not empty)
     Initialized an empty list ***L*’**,
     **For** each node ***id*** in ***L***
          Check each neighbor label ***i*** of ***n*’ *(id).list*** from ***N*’**,
          **For** each ***i***,
          1. Update the ***n*’*(i).len*** in ***N’*** to include the length from ***n(id).len***, and expand the label ***n*’*(id).id*** to ***n*’(*i*).*id***.
          2. Delete the ***id*** from neighboring list ***n*’(*i*).*list.*
          end**
          Attach all the neighboring label of n’ to ***L*’**,
      **end**
      Find only the unique label in ***L*’** to replace ***L***.
   **end**
   Find the maximum length value in ***N*’** and get the corresponding path from the label
**end**
~~~

### Calculation of persistence length, growth velocity and growth direction

To calculate the persistence length, *P*, we use the extracted backbone pattern. In short, the persistence length is a measure of how long distance the growing direction continues without curving. Since the persistence length can be measured with different models, and it is highly dependent on the ratio between the persistence length and contour length of the object, we implemented two different models. The first model is well suited when the object shows a high degree of curvature, that is when the persistence length is short in comparison to the contour length,

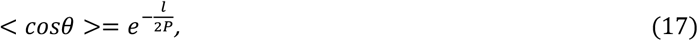

where a tangent-tangent correlation is measured using the angle *θ* between two points in the structure at a distance of *l*. Since we measure this in a two-dimensional surface, a 2 is added in the denominator (Lamour et al. 2014). The estimated persistence length is then measured by fitting this equation to the tangent-tangent correlation as a function of *l* ranging from the resolution of imaging system to the total length of the structure. For objects that are fairly straight, that is the persistence length is long in comparison to the contour length a more appropriate model is,

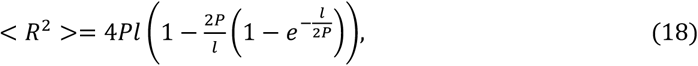

where the mean square of end-to-end distance between two points in the structure at each *l* is measured instead of the tangent-tangent correlation (Lamour et al. 2014).

To measure the growth velocity and growth directions we use the backbone pattern and measure differences between consecutive frames. The velocity is measured in the unit of area per time, thus it depends on the experimental settings which are related to the magnification of system, the size of the video recording area, number of pixels, and camera frame rate. The growth direction is given as degree of angles ranging from 0 to 360 where 0 indicates the direction along the horizontal line of the image to the right where the value increase for counter-clock wise changes.

## 6 Results and discussion

### Comparing Dseg with three algorithms and three Software

To validate the performance and accuracy of DSeg we compared the segmentation results of *S. coelicolor* hyphae with three commonly used segmentation functions as well as three well-established software. The segmentation functions were implemented in MATLAB and included; adaptive thresholding, the fast-marching level-set method (Sethian 1999), and the edge detection method with watershed transform (Meyer 1994). The tested software were B-COSFIRE (Strisciuglio and Petkov 2017), CellX (Dimopoulos et al. 2014) and TLM-Tracker (Klein et al. 2012).

Our validation shows that DSeg maintain the backbone pattern of the filamentous objects and distinguish the structure of objects better than the other tested methods (Fig S3A-G). We performed the comparison by using 100 frames of 512×512 pixels in resolution from a video that contain numerous *S. coelicolor* hyphae. We took frame 92 from the video as input to test the adaptive thresholding, the fast marching level-set, watershed, B-COSFIRE and CellX. The segmentation results of the obtained binary masks and are shown in Figure S3A-E. For methods A-D, the colored segmentation results is generated by labelling connect components using MATLABs function ‘*bwconncomp*’.

The three MATLAB functions were used for comparison and the binary masks extracted using the following approach. First, to get the segmentation result as binary mask for adaptive thresholding we used the MATLAB function ‘*adaptthresh*’. Second, the fast marching level-set method was based on the MATLAB function ‘*imsegfmm*’ with pre-selected seed locations marked as white dots presented in the color labeled image in Figure S3B. Third, the watershed transform method combines the MATLAB functions ‘*edge*’, ‘*Regional minima*’ and ‘ *watershed*’ to get object contours and to refine the results a size filter is applied.

To get the binary mask from the three software we used the following approach. Note that B-COSFIRE and CellX cannot perform tracking, therefore an image were used. For the B-COSFIRE method we used the default parameters (Fig S3D). For CellX, seed points were auto detected and the corresponding segmentation results for all seed points are shown in Fig S3E. To test dynamic growth from a video we assigned frames 83 to 92 to the video as inputs for TLM-Tracker and our DSeg software, in which the results in Fig S3F-G are shown for frame 92. The two color-labelled images in Fig S3F are segmentation results from TLM-Tracker software using the GAC level-set method and a rendering image from the results using MATLAB for comparison. The ground truth image marked manually containing not only backbone pattern but also branches.

Based on a ground truth of the segmentation result with *K* objects in total, we evaluated the score of segmentation for each method as follows:

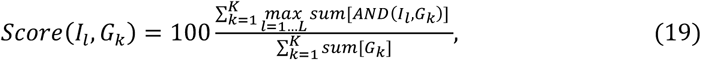

where *I*_*l*_ and *G*_*k*_ represents the binary image of segmentation results from one of the comparison methods and backbone pattern of ground truth respectively, *AND(A,B)* applies pixel by pixel logical and operation between two binary images *A* and *B, sum[A]* calculates the sum of values in *A* and *max* find the maximum value by checking all possible values, *l* and *k* represent the *l-th* object detected in *method* and *k*-th object in the ground truth, respectively. A perfect match results in a score of 100, and smaller value or 0 indicate less accurate segmentation results or results with large errors. To ensure fair comparison among results, in practice, the *I*_*l*_ is dilated with a 3 by 3 kernel because of human errors when drawing the ground truth. Note that to avoid evaluation error caused by one segmentation results covered in a large area to be evaluated multiple times, the *l-th* object selected from the *max* function can only be evaluated once. The score using frames in the sample video as well as the evaluation of the performance for all the comparison methods in this paper are presented in Table S1.

### Comparison of methods

Here we briefly discuss advantages and disadvantages of the compared methods to extract patterns from images. We first present the three MATLAB functions and thereafter three software. The *adaptive thresholding* method requires no pre-knowledge of the objects and can handle significant uneven intensity distribution in the image. However, due to its simplicity, this algorithm produces results with a lot of noise and irregular shapes making it very difficult to extract the skeleton pattern of objects later used for analysis. *Level-set* with *fast-marching* is a swift algorithm that can handle the topological changes of an object’s contour. The contours of the segmented object are smoother than that from the adaptive thresholding method, therefore it is easier for the morphological thinning operation to extract the skeleton structure. However, the results from this algorithm are dependent on the seed points selected by the user, and the segmentation areas are restricted to local regions. Finally, the *watershed approach* use image intensity and the gradient of intensity to get an object’s contour. An advantage with the watershed transform is that it can separate two different objects that are touching each other. However, the drawback is the over segmentation that makes merging of small fragments into an object very difficult.

We also tested three separate software. B-COSFIRE is a program designed to find line structures. We find that this program can find line structures much better than thresholding providing less noises and only a few irregular shapes. This algorithm does not produce over segmented fragments as seen when using the watershed transform. On the contrary, the results show clusters with several different objects which needs to be separated manually afterward the analysis. CellX is a powerful program for cell segmentation and optimized for objects with a clear membrane. We find that for objects with filamentous shape, the program cannot find the seed points correctly and thus the segmentation results are incorrect. TLM-Tracker is capable of tracking objects using time-lapse data only after objects have been segmented. Since the segmentation is an independent process the results from the software are equally good as using GAC algorithms alone.

To overcome the aforementioned drawbacks, we design our algorithm to be able to handle time-lapse data and to update the segmentation process using the history of detected objects. We use the binary level-set method combined with thresholding to make the algorithm fast, produce smoothed and objective object structure that fulfill local optimal condition according to equation 13, and suppress the influence from other objects in the background. In our method, the constraints in size and shape regularize the evolvement of the level-set function at each frame so that the errors caused by noises in intensity and filament structure are also limited. Therefore, object growth information extracted from the backbone pattern using our segmentation method is stable and more reliable comparing to other methods.

### Proof of concept using various static and video data

Figure S4 shows examples of how DSeg can quantify different parameters from the backbone structure of identified objects, for example, length, width and persistence length, using various types of static and time-serie images. The *Grow speed* of an object is measured by tracking the two ends of the backbone structure. The *Maximum angle* measures the largest angle between two tangents found along the objects curve. The *Orientation* is the angle between a vector directed horizontally to the right and a vector in the direction along the growing regions of an object’s long axis. This long axis is the major axis of an ellipse by fitting an elliptical shape to the object image.

**Table S1.** Evaluation of 7 segmentation methods using *S. coelicolor* videos of 474×492 pixels resolution and 32 frames recorded using DIC microscopy. Since CellX did not correctly detect seed points we could not include the accuracy value from this method. The notation + indicates that the time for the selection process is excluded for methods that does not assign seed point to the segmentation automatically. The notation ∼ is an estimate of the execution time. For TML-Tracker, we measured execution time of the GAC segmentation for 10 frames as video execution time and derive the averaged execution time for a single frame accordingly. The average performance of the DSeg using a single frame is 0.7s. Since the algorithm depends on the object number and structure complexity, we estimate the video execution time using 10 frames and 30 objects will take about 210 s to get the results in Figure S3G, excluding the time for selecting seed points.

| Method | Input | Seed | Detection | Detected Object # | Segmentation Accuracy (Score) | Frame Execution Time (s) | Video Execution Time (s) |
| --- | --- | --- | --- | --- | --- | --- | --- |
| **Adaptive thresholding** | Single image | No | No | 423 | 43.5 | 2.1 | - |
| **Fast-marching** | Single image | Yes | Semi-auto | 30 | 39.3 | 1.7+ | - |
| **Watershed** | Single image | No | No | 185 | 26.4 | 2.3 | - |
| **B-COSFIRE** | Single image | No | No | 112 | 63.5 | 1.6 | - |
| **CellX** | Single image | Yes | Auto | 74 | - | 15.9 | - |
| **TLM-Tracker** | Video data | Yes | Auto | 121 | 30.8 | 32 | ~320 |
| **Our method** | Video data | Yes | Semi-auto | 30 | 85.8 | 0.7 | ~210+ |

**Figure S3.**
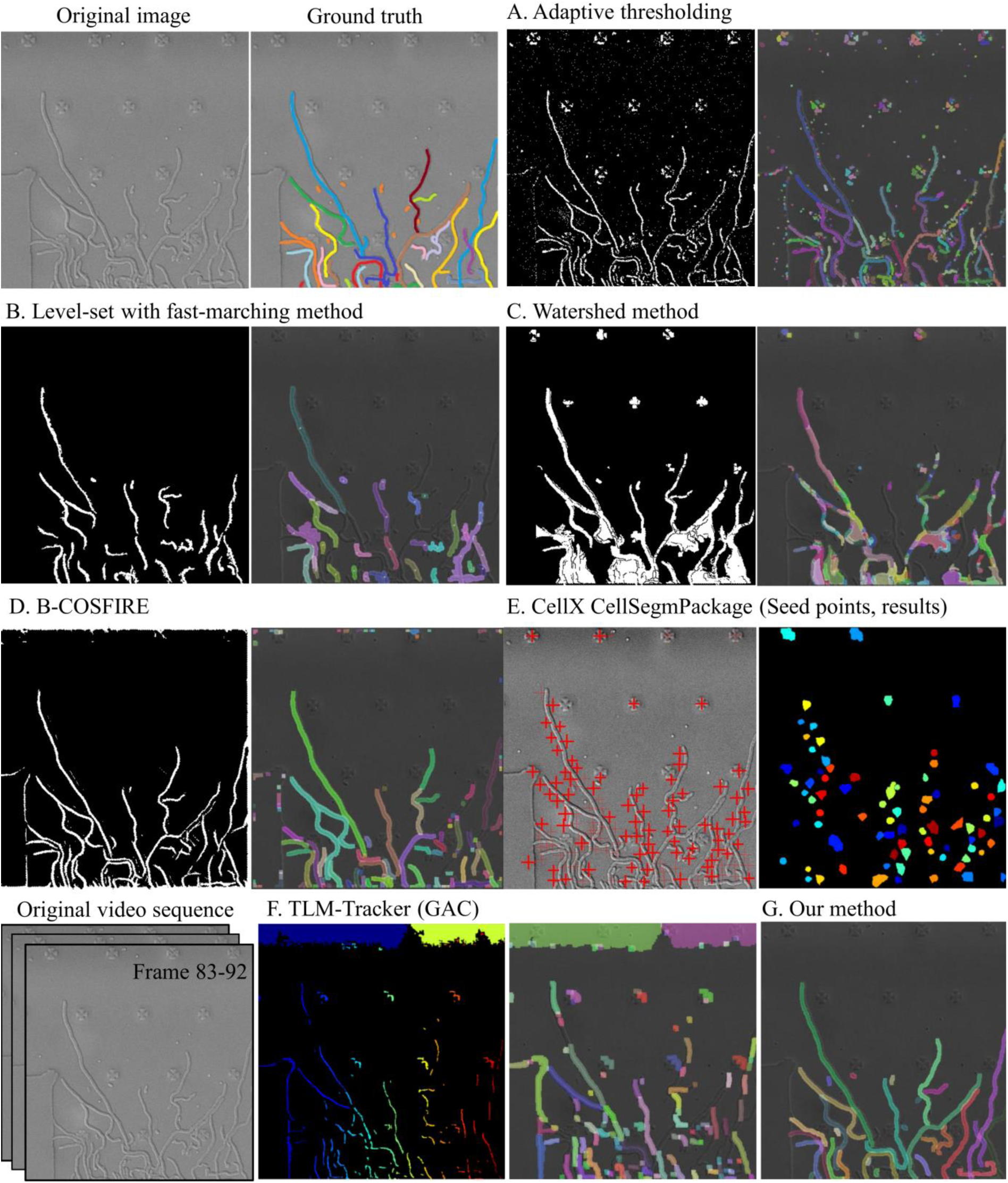
Segmentation results using *S. coelicolor* data under DIC microscopy. The original image taken from the frame 92 in the sample video is the input to method A-E. The video sequences extracted from frame 83-92 containing 10 frames are inputs for TLM-Tracker and our DSeg software labelled in F and G, respectively. The ground truth of segmentation results is marked manually based on the growth pattern analyzed from frame 1 to frame 92.

**Figure S4.**
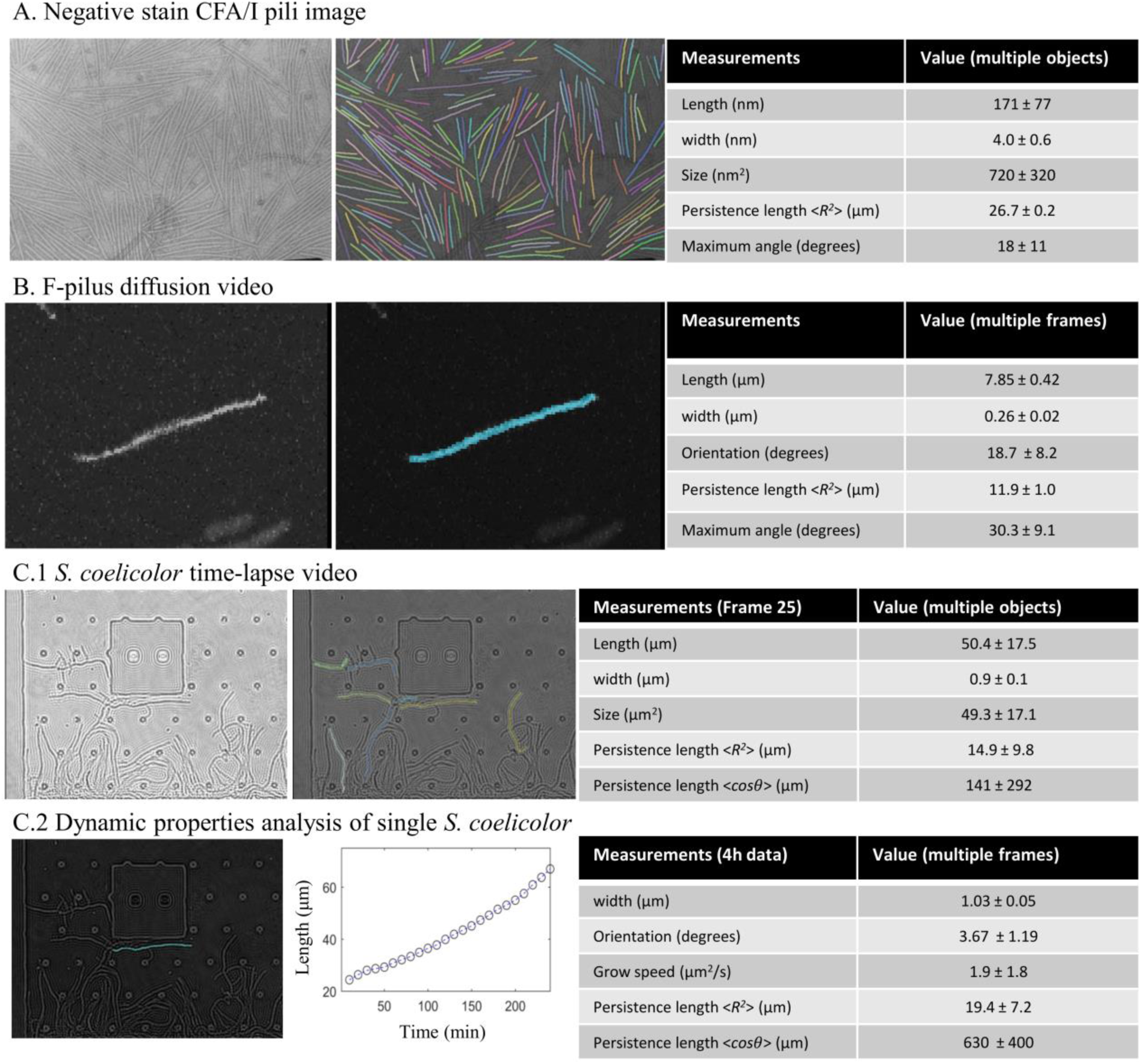
A) Negative stain data of CFA/I using transmission electron microscopy (left), segmentation results (middle), statistical dataset (right). The conversion factor is 0.423 nm/pixel (Brown et al. 2018). B) A video frame from a F-pilus diffusion video recorded using fluorescence microscopy (left), segmentation result (middle), statistical dataset (right). The video is taken from reference (Silverman and Clarke 2010). C.1) shows the static data analysis of *S. coelicolor* based on frame 25 in the video and C.2) presents the dynamic data analysis of single backbone structure. The *S. coelicolor* video are recorded using in-line digital holographic microscopy based on the setup in (Stangner et al. 2017; Zhang et al. 2017) with a conversion factor of 132 nm/pixel.

### Limitations of DSeg

Note that DSeg only corrects for linear translation, for example stage drift in the lateral plane. Therefore it is important to align the microscopy system with the sample carefully to avoid any rotation or tilt.

The software is design for tracking the shape of objects in 2D images. To obtain the object shape as correctly as possible, we recommend that the growth of any organism is restricted in 2D. This can be achieved using a thin flow-chamber device as demonstrated in our experimental procedure or by using an adhesive coated surface. 3D data cannot be analyzed directly but a slice of the 3D data can be segmented with the software.

The overlapping of objects cannot be handled completely by the software. However, in some cases when the time-lapse data contains growing objects covered partially by a small background object, the segmentation algorithm is capable of finding the backbone structure correctly. The segmentation result for data analysis only contains the backbone pattern of the object. The tracking of branches is not implemented in the software even though the shape of the branch is generated in the segmentation process.

